# Role of midbody remnant in meiosis II creating tethered polar bodies

**DOI:** 10.1101/191981

**Authors:** Alex McDougall, Celine Hebras, Gerard Pruliere, David Burgess, Vlad Costache, Remi Dumollard, Janet Chenevert

## Abstract

Polar body (PB) formation is an extreme form of unequal cell division that occurs in oocytes due to the eccentric position of the small meiotic spindle near the oocyte cortex. Prior to PB formation, a chromatin-centered process causes the cortex overlying the meiotic chromosomes to become polarized. This polarized cortical subdomain marks the site where a cortical protrusion or outpocket forms at the oocyte surface creating the future PBs. We observed that PB1 becomes tethered to the egg via PB2, indicating that the site of PB1 cytokinesis directed the precise site for PB2 emission. We therefore studied whether the midbody remnant left behind following PB1 emission was involved, together with the egg chromatin, in defining the precise cortical site for PB2 emission. During outpocketing of PB2 in ascidians, we discovered that a small corps around 1μm in diameter protruded from the center of the cortical outpocket that will form the future PB2, which we call the “polar corps”. During emission of PB2, this small polar corps became localized between PB2 and PB1 and appeared to link PB2 to PB1. We tested the hypothesis that this small polar corps on the surface of the forming PB2 was the midbody remnant from the previous round of PB1 cytokinesis. We had previously discovered that Plk1::Ven labeled midbody remnants in ascidian embryos. We therefore used Plk1::Ven to follow the dynamics of the PB1 midbody remnant during meiosis II. Plk1::Ven strongly labeled the small polar corps that formed on the surface of the cortical outpocket that created PB2. Following emission of PB2, this polar corps was rich in Plk1::Ven and linked PB2 to PB1. By labelling actin (with LifeAct::mCherry/GFP or TRITC-Phalloidin) we also demonstrated that actin accumulates at the midbody remnant and also forms a cortical cap around the midbody remnant in meiosis II that prefigured the precise site of cortical outpocketing during PB2 emission. Phalloidin staining of actin and immunolabelling of anti-phospho aPKC during meiosis II in eggs that had PB1 removed showed that the midbody remnant remained within the egg following emission of PB1. Dynamic imaging of microtubules labelled with Ens::3GFP, MAP7::GFP or EB3::3GFP showed that one pole of the second meiotic spindle was located near the midbody remnant while the other pole rotated away from the cortex during outpocketing. Finally, we report that failure of the second meiotic spindle to rotate can lead to the formation of two cortical outpockets at anaphase II, one above each set of chromatids. It is not known whether the midbody remnant of PB1 is involved in directing the precise location of PB2 in other species as in ascidians. However, a review of the literature indicates that PB1 is tethered to the egg surface via PB2 in a number of species including members of the cnidarians, lophotrochozoa and echinoids, suggesting that the midbody remnant formed during PB1 emission may be involved in directing the precise site of PB2 emission throughout the invertebrates.

## Introduction

Polar body (PB) emission occurs in oocytes and is an extreme form of unequal cell division. One defining feature of meiosis is that two successive rounds of M phase and cytokinesis occur without an intervening S phase. At the end of the first meiotic M phase (meiosis I) in animal oocytes, cytokinesis creates PB1 while at the end of the second meiotic M phase (meiosis II) cytokinesis produces PB2. This generates one oocyte and two polar bodies (although 3 PBs are sometimes observed since PB1 can divide). In chordates, this extreme form of unequal cell division depends on the actin-dependent migration of the first meiotic spindle from the oocyte interior to the oocyte cortex (Azoury et al., 2008; Prodon et al., 2006; Schuh and Ellenberg, 2008). Once at the cortex, the chromosomes cause a small subdomain of the overlying cortex to become polarized (Eager et al., 1976; Maro et al., 1984; Maro et al., 1986). More recent findings in mouse oocytes have demonstrated that a chromatin-centered Ran-GTP gradient polarizes a cortical subdomain in close proximity to the meiotic chromosomes (Deng et al., 2007). In mouse oocytes, the chromatin-centered Ran-GTP gradient has been observed using a Fret-based biosensor (Dumont et al., 2007). These findings were prompted by earlier work showing that spindle assembly can be caused by a chromatin-centered Ran-GTP gradient induced by chromatin localized RCC1 (Heald et al., 1996; Karsenti and Vernos, 2001; Karsenti et al., 1984). One essential difference therefore between PB formation and mitotic cytokinesis is that a chromatin-centered Ran-GTP gradient causes a subdomain of the cortex adjacent to the meiotic chromosomes to become polarized driving PB outpocketing (Dehapiot et al., 2013; Deng et al., 2007).

PB formation is a stepwise process beginning with cortical polarization, followed by protrusive outpocketing of the polarized cortex and ending with constriction of the cortical outpocket. During meiosis I in mouse oocytes, a chromatin-centered Ran-GTP gradient promotes the formation of an actin cap via the inactivation of ERM (Ezrin/Radixin/Moesin) independent of Cdc42 (Dehapiot and Halet, 2013). Cortical outpocketing is initiated at the actin cap during anaphase and is accompanied by the creation of dynamic actin via the recruitment of Cdc42, N-WASP and Arp2/3 in mouse (Dehapiot et al., 2013) and *Xenopus* oocytes (Ma et al., 2006; Zhang et al., 2008). Cdc42 is required for cortical outpocketing in mouse oocytes, since the dominant negative Cdc42 or specific deletion of Cdc42 prevents cortical outpocketing (Dehapiot et al., 2013; Wang et al., 2013). It is not entirely known how the fall in MPF activity triggers the cortical recruitment of active Cdc42 during anaphase, which can be abolished by preventing the fall in MPF activity (Zhang et al., 2008). Outpocketing also occurs during meiosis II, and similarly to PB1 emission, in mouse oocytes the active form of Cdc42 becomes enriched at a cortical subdomain adjacent to one spindle pole and chromatids that are closest to the cortex (Dehapiot et al., 2013). During PB cytokinesis, ECT2 localized at the spindle midzone leads to the formation of a RhoA contractile ring rich in myosin II (see review by Maddox et al., 2012). Thus, cortical outpocketing during anaphase I and II is thought to be induced by the cortical accumulation of active Cdc42 in *Xenopus* and mouse oocytes. However, it is not entirely clear what triggers the accumulation of active Cdc42 at the cortex driving outpocketing (Leblanc et al., 2011). Here in the ascidian we have discovered that during meiosis II the midbody formed at PB1 emission participates in directing the precise site of PB2 outpocketing. This results in PB2 being emitted at the precise site of PB1 cytokinesis, thereby resulting in PB1 becoming tethered to the egg indirectly via PB2. Such tethered polar bodies appears to be a widespread occurrence throughout invertebrate species including cnidarians, lophotrochozoa, and echinoderms.

## Results

Ascidian oocytes are arrested at metaphase of meiosis I, and they extrude both PBs within a half hour of fertilization (McDougall and Sardet, 1995). These two PBs are always emitted at precisely the same site of the egg surface. Careful time-lapse observations showed that a small polar corps formed on the cortical surface of a protruding outpocket that will form PB2, and also that this small polar corps linked PB1 to PB2 (Figure 1Ai and Supp Movie 1). As a consequence of this precise spatial control, PB1 became tethered to the egg indirectly via PB2 (see Figure 7 scenario 1). To determine the precise location of the polar corps on the cortical outpocket, we analyzed the angle between the polar corps and the apex of the outpocket (Figure 1Aii). The polar corps was centered within 9° of the cortical outpocket apex (81.4°+/- 1.4°, mean +/- sem., n=15 and Figure 1 Aii). We also show that PB1 is tethered to the egg surface via PB2 in the bivalve *Mytilus galloprovincialis* (Figure 1B and Supp Movie 2). Indeed, published images of oocytes and their polar bodies from a number of invertebrate species show that PB1 is almost always tethered indirectly to the egg surface via PB2 (see Figure 7 model and Table 1). In order to be tethered to each other, PB2 must be extruded almost exactly at the previous site of PB1 formation – delocalization by as little as about half the length of the second meiotic spindle would abolish the tethering between polar bodies and instead PB1 and PB2 would each be linked to the egg surface independently (see model, scenario 2: Figure 7).

**Table 1.** Tethered polar bodies. A number of species display tethered first and second polar bodies as in the ascidian and depicted in Scenario 1. This is not an exhaustive list since we have noted several other examples that are not detailed here. Notable exceptions are the vertebrates that do not show tethered polar bodies. *Center for Cell Dynamics website: http://rusty.fhl.washington.edu/celldynamics/gallery/index.html

| **Species** | **PBs tethered** | **Publication** |
| --- | --- | --- |
| **Jellyfish**<br>*Clytia hemispherica* | **Yes** | (Amiel et al., 2009) |
| **Nemertean worms**<br>*Micura alaskensis* | **Yes** | (Maslakova, 2010) |
| **Sea slug**<br>*Cuthona lagunae* | **Yes** | (Goddard, 1991) |
| **Clam**<br>*Acila castrensis* | **Yes** | Von Dassow<br>Center for Cell<br>Dynamics website* |
| **Mussel**<br>*Mytilus galloprovincialis* | **Yes** | This study |
| **Starfish**<br>*Asterina pectinifera* | **Yes** | (Harada et al., 2003) |
| **Sea cucumber**<br>*Holothuria moebi* | **Yes** | (Miyazaki et al., 2005) |
| **Ascidian**<br>*Phallusia mammillata* | **Yes** | This study |
| **Nematode**<br>*Caenorhabditis elegans* | **Possibly** | (Dorn et al., 2010) |
| **Xenopus** | **No** | (Li et al., 2016) |
| **Mouse** | **No** | (Dehapiot et al., 2013) |

**Figure 1.**
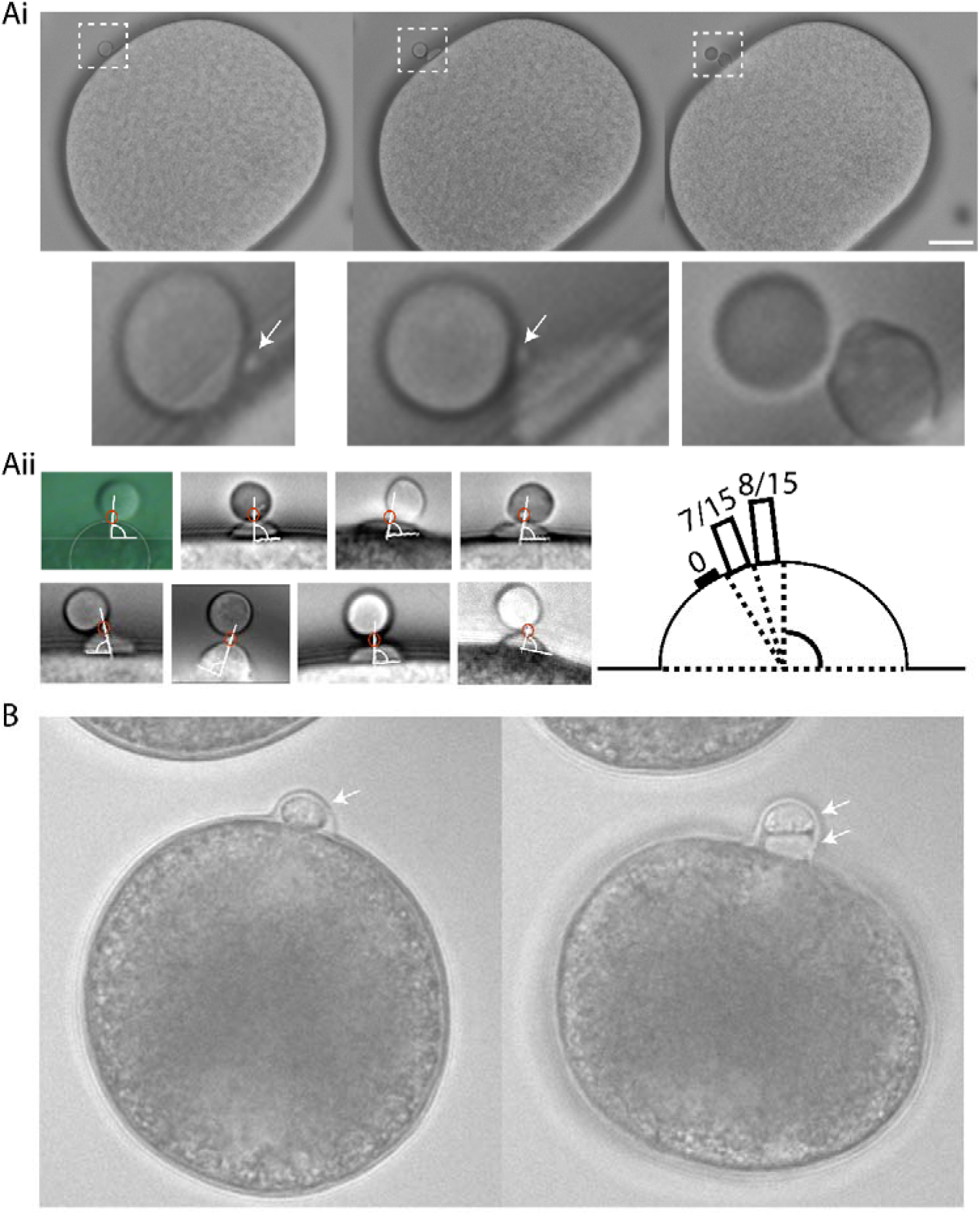
A small protrusion forms on the surface of the PB2 outpocket in *Phallusia*. A i) Bright field images from a time-lapse experiment of fertilized *Phallusia mammillata* eggs. During polar body 2 (PB2) emission, a small polar corps can be observed on the surface of PB2 outpocket (boxed region). Enlarged views of the boxed regions at the bottom show in greater detail the small protrusion which is present before outpocketing begins (bottom left, arrow) and remains present during outpocketing (bottom middle, arrow). n>50 eggs. Scale bar = 20μm. See Supp Movie 1. ii) Analysis of angle between polar corps (red circle) and outpocket center. Diagram illustrating a summary of the polar corps position at 61-70°, 71-80° and 81-90° (number of eggs in brackets). Mean polar corps position was 81.4°+/-1.4° +/-sem, n=15. B) Bright field images from a time-lapse experiment showing PB2 emission under PB1 in *Mytilus galloprovincialis.* PB1 is indicated by the arrow in the first image, and both PB1 and PB2 by the two arrows in the second image. n>50. See Supp Movie 2.

We sought to determine whether this small polar corps was the midbody remnant from the first meiotic division using live imaging of Polo kinase (Plk1). Plk1 is a conserved component of the spindle midzone and midbody remnant in many cell types (Hornick et al., 2010; Li et al., 2010), including mouse oocyte where it localizes to the first midbody formed during PB1 emission (Sun et al., 2012; Wianny et al., 1998). We had previously found that ascidian Plk1::Ven strongly labels the central spindle and midbodies in mitotic cells of the ascidian embryo (McDougall et al., 2015). Using live fluorescence imaging of Plk1::Ven, we followed more precisely the location of the midbody following PB1 emission (Figure 2A). The small polar corps labelled with Plk1::Ven and was present on the surface of the egg next to the site where PB1 was attached to the egg throughout meiosis II (Figure 2A, Egg1). During PB2 emission, this small polar corps labelled with Plk1::Ven protruded from the cortical outpocket of the forming PB2 and eventually linked PB2 to PB1 (Figure 2A, arrows in Eggs 2, 3 and 4). The meiotic spindle and the midbody were labelled simultaneously by co-injecting EB3::3GFP to label microtubules together with Plk1::Rfp1 mRNA (Figure 2.B). Plk1::Rfp1 again labelled the midbody that formed at the apex of PB2 outpocket (Figure 2B). Note also that Plk1::Rfp1 also labelled the chromosomes in Meta I (Figure 2B). Finally, a confocal z-stack through a live fertilized egg clearly shows the position of midbody 1 between PB1 and PB2, and midbody 2 between PB2 and the egg (Figure 2C and Supp Movie 3). Plk1::Ven also labeled the midzone and the midbody in the embryo (Figure 2D).

**Figure 2.**
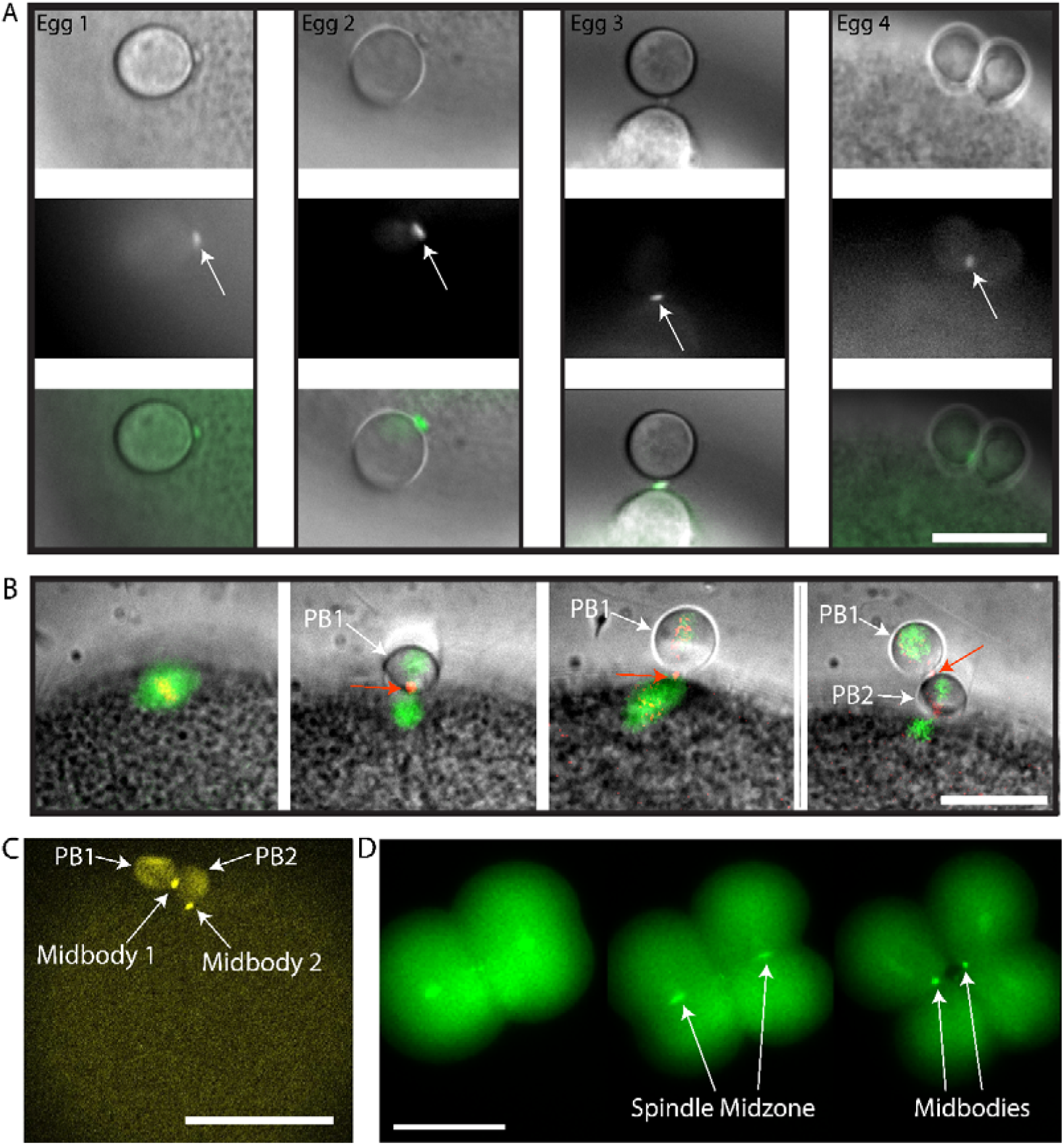
Plk1::Venus labels the midbody that forms between PB1 and the egg. A) Four different examples of the small polar corps labelled with Plk1::Ven. Upper row of images shows bright field images of PB2 emission site adjacent to PB1. Middle row of images shows that Plk1::Ven labelled the midbody that formed between PB1 and the egg (see arrows). Bottom row is the overlay. Plk1::Ven localization to the midbody remained during the process of PB2 emission (see arrows). n=12. Scale bar = 20μm. B) Epifluorescence images of meiotic spindle labelled with EB3::3GFP and the midbody with Plk1::Rfp1. Note that Plk1::Rfp1 labels the chromosomes (red) on the Meta I spindle (first image), then the midbody (second image) and also the midbody is found at the apex of the PB2 outpocket (third image). PB1 is tethered to PB2 (fourth image). First midbody is indicated with red arrows and PBs are indicated by the white arrows. Scale bar = 20μm. n=5. C) Fertilized egg after PB1 and PB2 emission. Confocal image from a z-stack showing the localization of Plk1::Ven to midbody 1 and midbody 2 (arrows). PB1 and PB2 are also indicated by arrows. n=12. Scale bar = 50μm. See Supp Movie 3. D) Two to four cell-stage. Epifluorescence images of midbody formation. Plk1::Ven labels the central spindle (arrows) then both midbodies (arrows) at the end of cytokinesis. n= 22. Scale bar = 50μm.

In order to determine whether the midbody remnant of PB1 was located in the egg or PB1, we removed PB1 by gentle pipetting and searched for labelling of the midbody remnant in such PB1-free eggs. First we used LifeAct::mCherry to label actin in live fertilized eggs and show that a cortical actin cap forms under PB1 around the PB1 midbody during meiosis II in the ascidian, and prefigured the cortical site where outpocketing occurred (Figure 3A, Supp Movies 8 and 9). Next we pipetted fertilized eggs during meiosis II to remove PB1, fixed and performed Phalloidin staining to observe the actin accumulated at the PB1 midbody. In eggs that had their PB1 removed, the PB1 midbody was still present in the egg (Figure 3B). Also note that although the egg chromosomes are located near the midbody they are not always located directly under the midbody (Figure 3Biii). In addition, we have found recently that anti-phospho aPKC (atypical protein kinase C) strongly labels the midbody (Pruliere et al., in preparation). We thus also examined eggs that had PB1 removed with anti-phospho aPKC to label the midbody, anti-tubulin for microtubules and DAPI for DNA. The midbody remnant was again located in the egg following removal of PB1 (Figure 3C). We thus conclude that the first midbody remains within the egg following PB1 emission.

**Figure 3.**
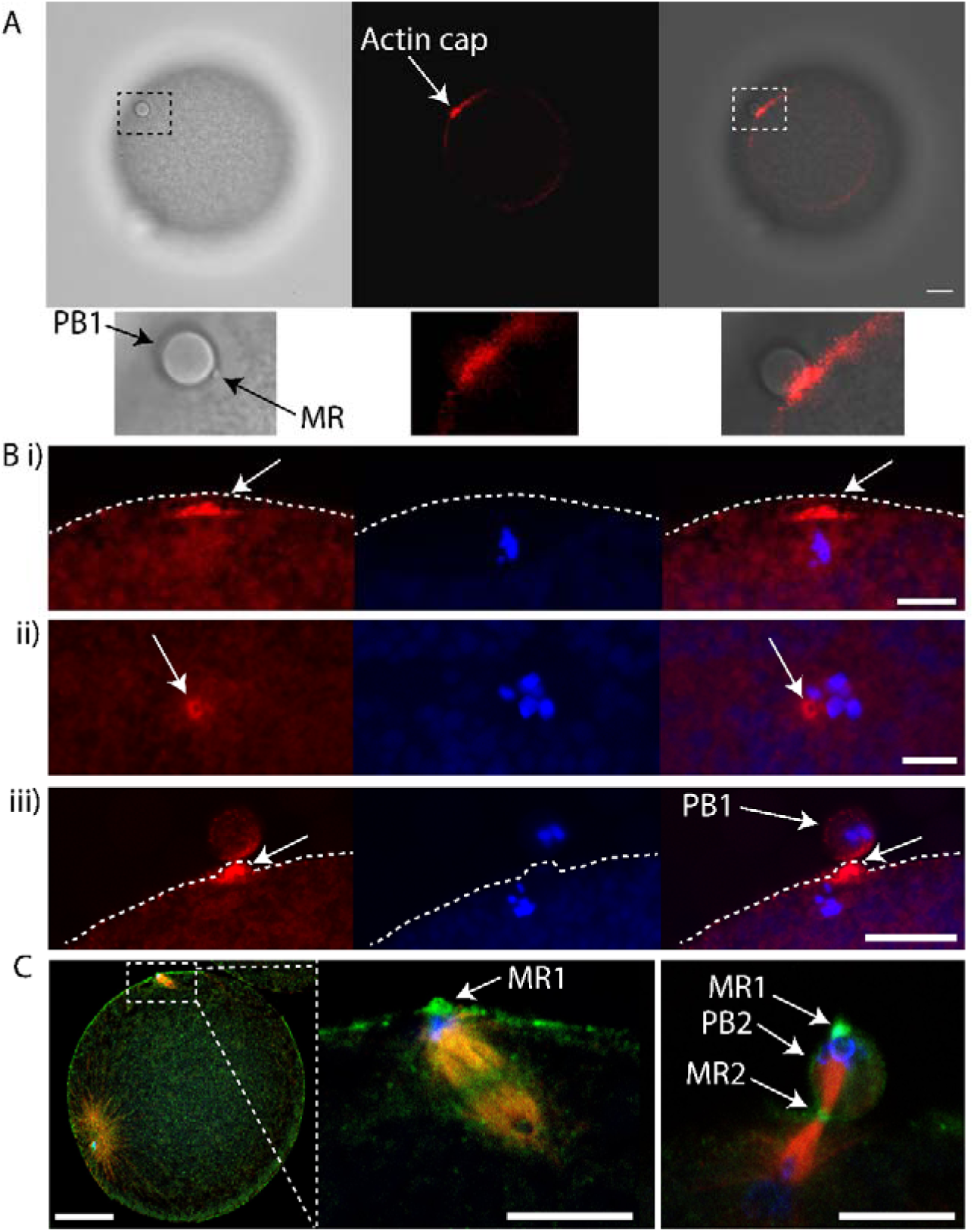
First midbody remains in the egg following PB1 emission. A) Unfertilized eggs were injected with LifeAct::mCherry protein to label actin and fertilized. Confocal images showing actin cap and midbody remnant (MR) indicated by arrows. Left, bright field, middle LifeAct, right overlay. Insets of boxed region at bottom. Scale bar = 20μm. n=8. See Supp Movies 8 and 9. B) Fertilized eggs were fixed during meiosis II and stained using Phalloidin::TRITC to label actin and Hoechst to label chromosomes. i) Confocal images showing actin cap (arrows) during meiosis II in an egg following PB1 removal. ii) Confocal images showing actin labelling of the midbody (arrows) during meiosis II in an egg following PB1 removal (arrows). iii) Control egg that displayed PB1 attached to the egg surface. Note that the midbody is strongly labelled with Phalloidin (arrows). Dotted line indicates surface of the egg. Scale bars = 10μm. n>50. C) Fertilized eggs were pipetted during meiosis II to remove PB1, fixed and labelled with anti-phospho aPKC (green), anti-tubulin (red) and DAPI (blue). Left image: overlay showing the tilted second meiotic spindle. The sperm aster is also visible, far left. Scale bar=30μm. Middle image: inset of boxed region showing that one pole of second meiotic spindle is aligned with PB1 midbody remnant (MR1 arrow). Note that PB1 was removed by pipetting. Scale bar=10μm. Right image: Another egg which had already emitted PB2 showing location of first midbody remnant (MR1, arrows), second midbody remnant (MR2, arrow) and PB2 (arrow). Note that PB1 was removed by pipetting. Scale bar=10μm

Next we wished to monitor the behavior of the second meiotic spindle in order to determine its dynamics during PB2 emission. In order to monitor microtubules of the second meiotic spindle, we microinjected eggs with mRNA encoding Ens::3GFP mRNA (we also used MAP7::GFP or EB3::3GFP) and incubated them overnight to allow translation of the fluorescent protein (McDougall et al., 2015). We noted that the second meiotic spindle tilted during emission of PB2, with one pole anchored near the site where PB2 was emitted (Figure 4 and Supp Movie 4). Initially the spindle was centered under PB1, then it moved laterally along the cortex and tilted (Figure 4 and Supp Movie 4). We therefore propose that the remnant of the PB1 midbody and the subdomain of cortex polarized around the midbody attracts one pole of the second meiotic spindle, causing one pole of the spindle to enter the cortical outpocket and tilt during PB2 emission.

**Figure 4.**
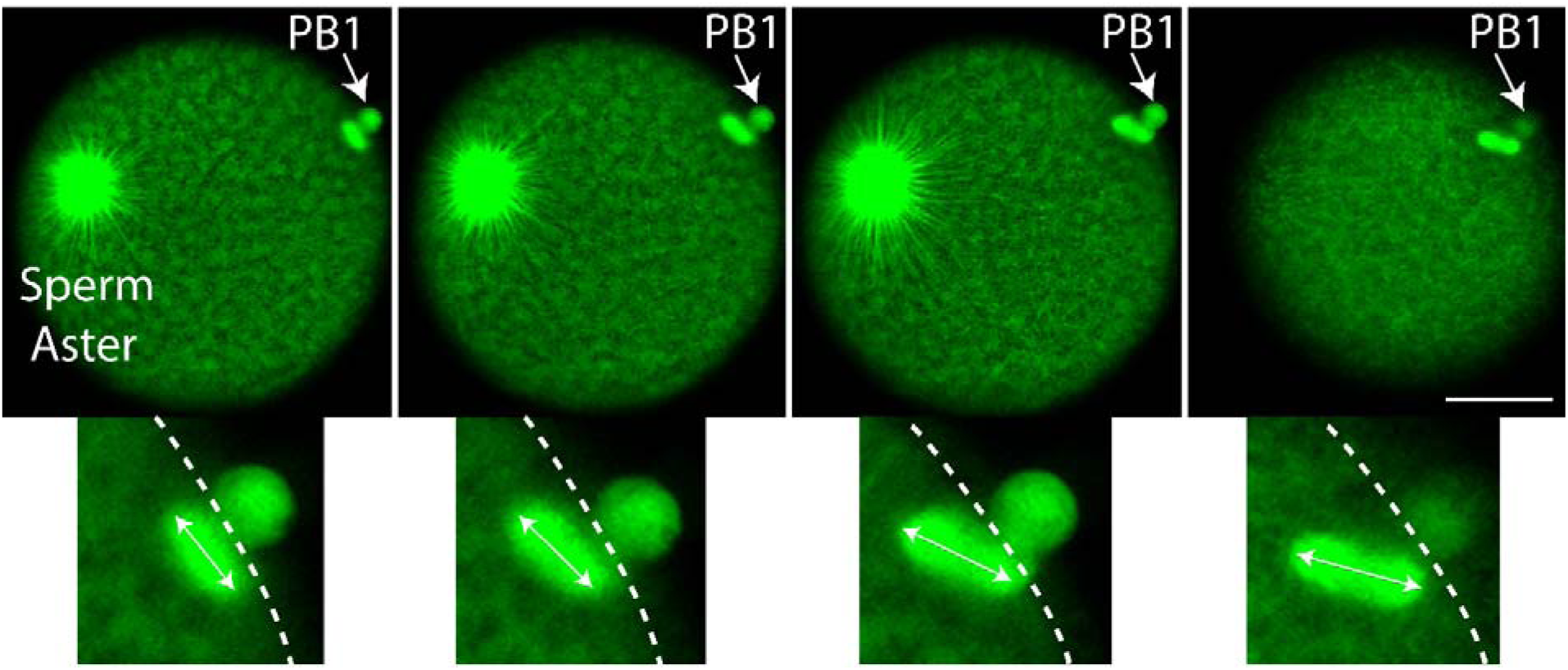
Second meiotic spindle tilts during PB2 emission. Eggs were previously microinjected with mRNA encoding Ens::3GFP to label the microtubules (green). Confocal images extracted from a time-lapse experiment showing the tilt of the second meiotic spindle. Upper row of images showing that the second meiotic spindle lies under PB1 parallel to the cortical surface, then begins to tilt (image 2) and continues to tilt (images 3 and 4) as PB2 is emitted (see last image of Supp Movie 4). Insets at the bottom show more clearly the tilt of the second meiotic spindle (double headed arrow shows spindle orientation). Large sperm aster is also visible. n=12. Scale bar = 40μm. Time interval between images is 1 min. See Supp Movie 4.

We sought to determine whether chromatin could cause polarization of the cortex in the ascidian as in the mouse (Deng et al., 2007), and also determine the consequences of chromatin-induced polarization of the cortex when the spindle failed to tilt. We have observed that two cortical outpockets, one on either side of PB1, sometimes occur during failed PB2 emission (see Supp Movie 5). Careful observation revealed that, on occasion, the second meiotic spindle failed to rotate. Two patches of chromatin accumulated at the spindle poles as during normal anaphase I (Figure 5A and Supp Movie 6). However, and more importantly, the cortex formed protrusive outpockets above both sets of chromatids and spindle poles leading to the emission of two simultaneous PB2 outpockets (Figure 5A and Supp Movie 6). Finally, when spindle microtubules are depolymerized with nocodazole, cortical outpocketing still occurred next to the chromosomes (Figure 6 and Supp. Movie 7) showing that chromatin proximity is sufficient to drive cortical outpocketing in the ascidian.

**Figure 5.**
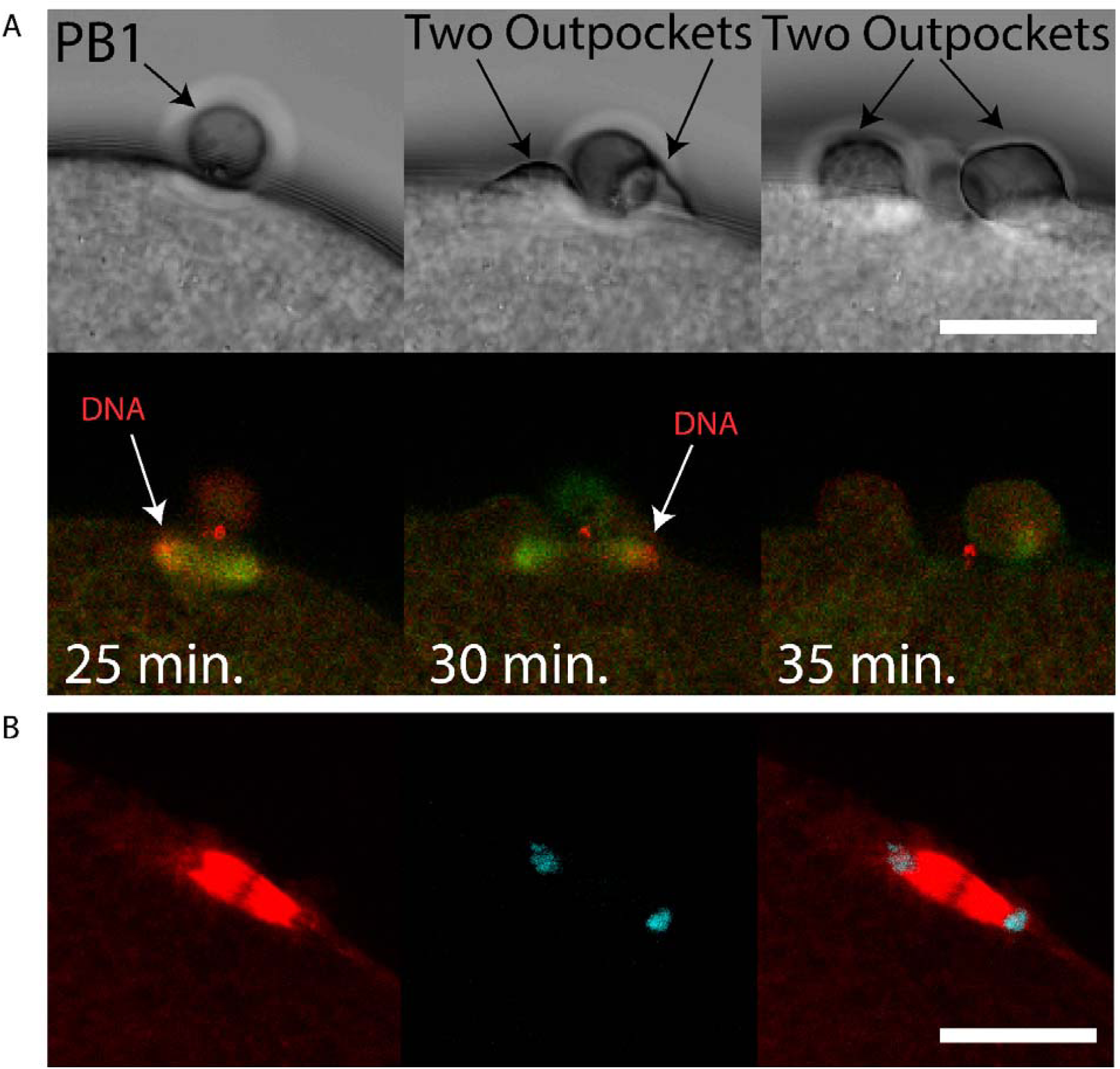
Failed rotation of second meiotic spindle giving two PB2 outpockets. A) Unfertilized eggs were injected with mRNAs encoding Ens::3GFP (microtubules green) and Kif2::mCherry (chromosomes red) mRNA. Two outpockets are shown in the second and third bright field images respectively (arrows). Note that the second meiotic spindle fails to rotate and two outpockets form above both sets of chromosomes following Ana II (see 24-31 min. on Supp Movie 6). Note also that Kif2::mCherry also labels the midbody (prominent red staining near PB1). Scale bar = 20μm. n=5. See Supp Movie 6 for the full data-set. Another example of two simultaneous PB2 outpockets is shown in Supp Movie 5. B) Fertilized eggs were fixed and labelled with anti-Tubulin (red) and stained with DAPI (blue). Confocal images of a spindle positioned parallel to the egg cortex in anaphase. Scale bar = 20μm, n=5.

**Figure 6.**
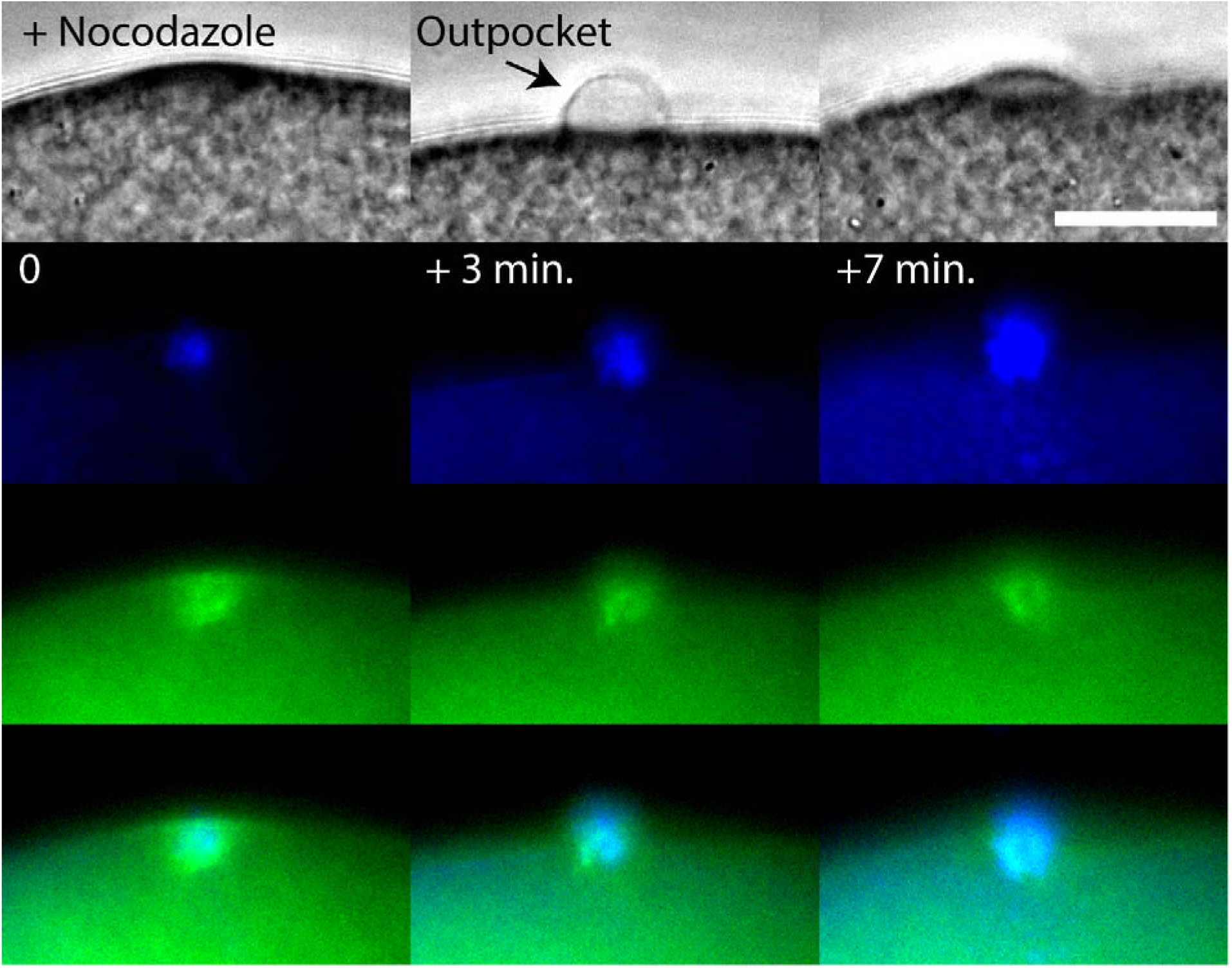
DNA can polarize the egg cortex in ascidians as in mouse oocytes. Unfertilized eggs were injected with mRNA encoding MAP7::GFP to label the microtubules. The following day the injected eggs were bathed in Hoechst 33342 (10μg/ml) to label the chromosomes and 2μM nocodazole to depolymerize the microtubules and then fertilized. Upper row of images: bright field; second row of images: chromosomes blue; third row of images: microtubule green; bottom row of images: overlay of green and blue images. Time between images is displayed on the second row of images. The meiotic spindle microtubules depolymerized and formed a depolymerized mass around the egg chromosomes (first column of images). A cortical outpocket above the DNA forms (second column of images). The outpocket resorbs into the egg (last column of images). Scale bar = 20μm. n=8. See Supp Movie 7.

In this report we show that PB2 is emitted at the site of PB1 cytokinesis. Our results show that the midbody formed during PB1 cytokinesis sits at the center of the cortical outpocket that protrudes from the egg surface during PB2 emission, and that following PB2 emission the first midbody linked PB1 to PB2. PB1 therefore became tethered indirectly to the egg via PB2. In addition, we show that during PB2 cortical outpocketing, one pole of the second meiotic spindle enters the cortical outpocket accompanied by rotation of the second meiotic spindle. Finally, we demonstrate that failure of the second meiotic spindle to rotate can lead to two simultaneous cortical outpockets rather than emission of one PB2.

## Materials and Methods

### Origin of the animals

Adult animals of *Phallusia mammillata* and *Mytilus galloprovincialis* were collected at Sète (Etang de Tau, Mediterranean coast, France). Ascidian gamete collection, dechorionation, fertilization and embryo cultures were as described previously (Sardet et al., 2011).

### Microinjection, imaging and reagents

Microinjection was performed as previously described (McDougall et al., 2014). Briefly, dechorionated eggs were mounted in glass wedges and injected with mRNA (1-2 μg/μl pipette concentration/ ~1-2% injection volume) using a high pressure system (Narishige IM300). mRNA-injected eggs were left for 2-5 hours or overnight before fertilization and imaging of fluorescent fusion protein constructs. Epifluorescence imaging was performed with an Olympus IX70, Zeiss Axiovert 100 or Axiovert 200 equipped with cooled CCD cameras and controlled with MetaMorph software package. Confocal microscopy was performed using a Leica SP5 or SP8 fitted with 40x/1.3na oil objective lens and 40x/1.1na water objective lens. All live imaging experiments were performed at 18-19°C. Nocodazole was prepared as a stock solution at 10 mM in DMSO and used at a final concentration of 2 μM. Hoechst 33342 was prepared as a stock solution of 10 mg/ml in DMSO and used at 10 μg/ml to label chromosomes.

Fixation and labelling for immunofluorescence. Eggs were fixed in -20° methanol containing 50 mM EGTA, blocked with PBS containing 2% BSA, and incubated with anti-tubulin primary antibody DM1a (Sigma-Aldrich) at a dilution of 1:500 and TRITC-conjugated anti-mouse secondary antibody (Santa Cruz) at a dilution of 1:200, washed in PBS and mounted in Citifluor (Biovalley, AF1-100). Using the same fixation procedure, fixed eggs were incubated with anti-phospho aPKC antibody (Santa Cruz, sc-12894-R) at a dilution of 1:100 and FITC-conjugated anti-rabbit secondary antibody (Santa Cruz) at a dilution of 1:200. For TRITC-Phalloidin (Molecular Probes, R415), activated eggs were fixed in 3.7% formaldehyde in 0.5 M NaCl in PBS 10 and 12 min. after PB1 emission. After several washes in PBT (PBS containing 3% BSA and 0.05% Triton X-100), the eggs were stained using TRITC-Phalloidin (10 μg/ml) and Hoechst 33342 (1 μg/ml). LifeAct::GFP or LifeAct::mCherry protein was made in bacteria, purified (8 μg/μl) and injected into unfertilized eggs.

### Synthesis of RNAs

We used the Gateway system (Invitrogen) to prepare N and C-terminal fusion constructs using pSPE3::RFA::Venus, pSPE3::Venus::RFA and pSPE3::RFA::Rfp1 (a gift from P. Lemaire), plus pSPE3::Rfp1::RFA, pSPE3::RFA::mCherry and pSPE3::mCherry::RFA. All synthetic mRNAs were transcribed and capped with mMessage mMachine kit (Ambion). Gene models and origin of all GFP-type constructs used (Ens::3GFP, Plk1::Ven, MAP7::GFP, EB3::3GFP) can be found in our methods article (McDougall et al., 2015), and Kif2 constructs in Costache et al. Nat. Comm. (in press). Briefly, synthetic mRNAs for the various constructs (Plk1::Ven, Ens::3GFP, MAP7::GFP, Kif2::mCherry) were microinjected into unfertilized *Phallusia* eggs which were left overnight to translate fluorescent fusion protein products (McDougall et al., 2015).

## Discussion

Although a Ran-GTP gradient emanating from the chromosomes of the meiotic spindle causes the overlying cortex to polarize in readiness for PB formation (Dehapiot and Halet, 2013; Deng et al., 2007), we propose here in ascidians that the precise location of PB2 is dictated by the midbody remnant left behind in the egg following emission of PB1. We show that the midbody remnant formed during PB1 formation remains in the egg following PB1 emission and sits at the apex of the cortical outpocket that will form PB2. This situation is similar to the finding in somatic cells where midbody remnants remain in one of the two daughter cells (Chen et al., 2013; Guizetti and Gerlich, 2010) rather than being externalized following cell division (Crowell et al., 2014). Once PB2 has been emitted, the midbody remnant therefore links PB1 to PB2, and thus PB1 becomes indirectly tethered to the egg surface via PB2 (see Figure 7, scenario 1). We thus use the phrase “tethered polar bodies” to reflect scenario 1 in Figure 7 whereby PB1 is tethered to the egg indirectly via PB2 (Figure 7). Note that not all species display tethered polar bodies, and instead that both PB1 and PB2 can be linked to the egg surface directly (Figure 7, scenario 2). Due to the widespread occurrence of tethered polar bodies within the invertebrates including ascidians, we thus came to test the hypothesis that the midbody formed during emission of PB1 may direct the precise site of PB2 formation.

**Figure 7.**
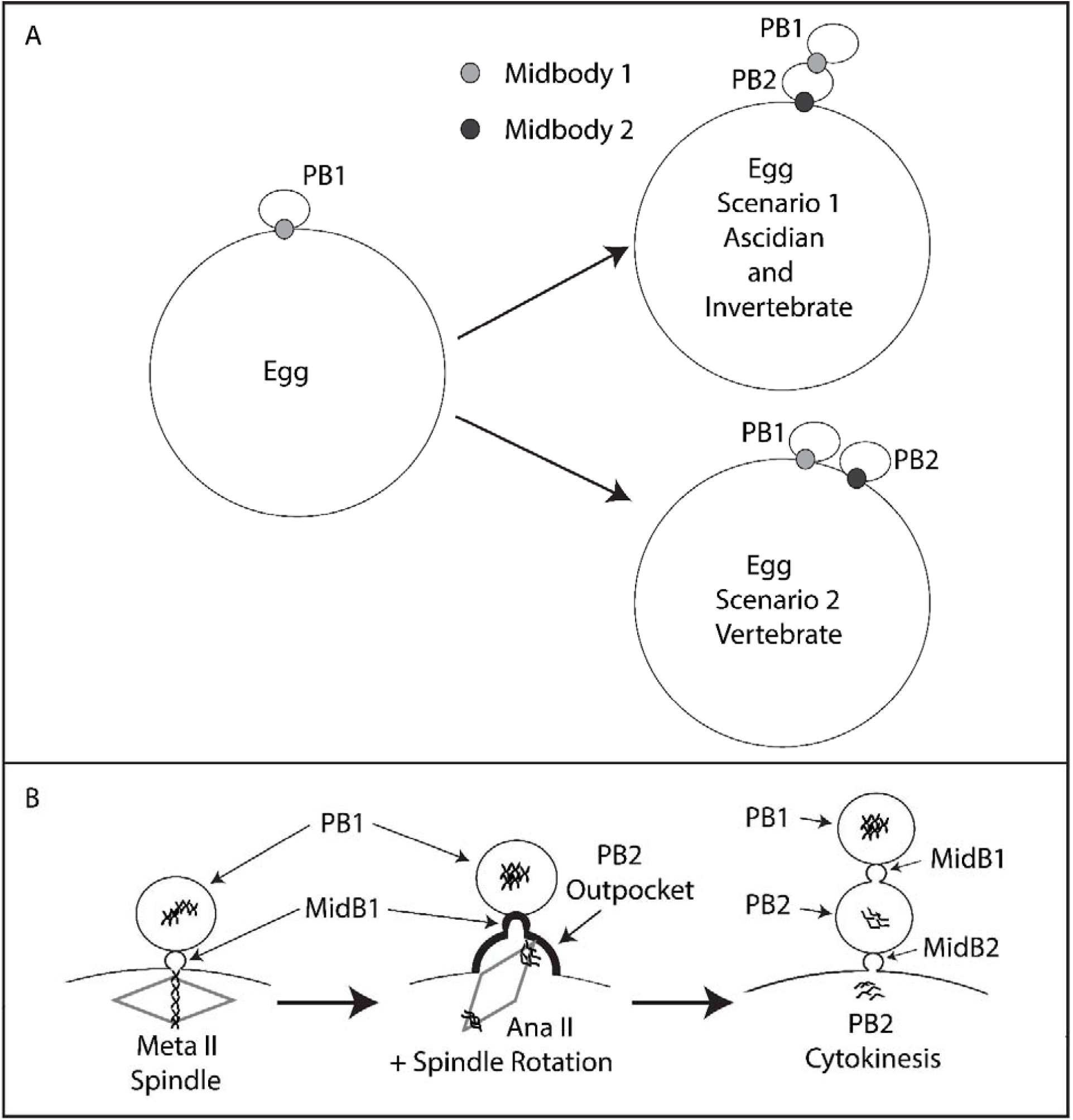
Model. Tethered polar bodies. A) Scenario 1: tethered polar bodies. Following emission of both polar bodies, the first midbody attaches PB1 to PB2 while the second midbody links PB2 to the egg. PB1 is thus tethered to the egg indirectly via PB2. Scenario 1 represents ascidians and maybe many invertebrates (see Table 1 for details). Scenario 2: the first midbody links PB1 to the egg and the second midbody also links PB2 to the egg. Scenario 2 represents mouse and *Xenopus*. Midbody 1 is depicted as light grey, midbody 2 as dark grey. However, it should be stressed that midbody position is only known for certainty in the ascidian. B) PB2 emission dynamics. Midbody 1 is located at the apex of PB2 cortical outpocket. Spindle rotation into PB2 cortical outpocket is displayed. PB2 is emitted attached to PB1 via midbody 1 (MidB1) and PB1 is thus tethered indirectly to the egg via PB2.

An analysis of published images of eggs with two polar bodies indicates that a diverse array of species show that PB1 is tethered to PB2 instead of being linked directly to the surface of the egg (Figure 7, scenario 1). For example, in jellyfish PB1 is tethered to PB2 instead of being linked directly the egg surface (Amiel et al., 2009). Even clearer examples are offered by species belonging to the lophotrochozoa. First, in the nemertean *Micura alaskensis* PB1 is tethered to the egg via PB2 (Maslakova, 2010). Likewise, in the nudibranch *Cuthona lagunae* PB1 is tethered to the egg via PB2 (Goddard, 1991). In marine bivalves, PB1 and PB2 are again tethered, for example in *Acila castrensis* (see Von Dassow, Center for Dynamics and Table 1 for link to website). We also noted that PB1 and PB2 are tethered in our time-lapse recordings of polar body emission in the mussel *Mytilus galloprovincialis* (Figure 1B and Supp. Movie 2). Amongst the ecdysozoa it is less clear whether PB1 is tethered to PB2. For example, in shrimp oocytes PB1 is located outside the hatching envelope far from PB2 (Hertzler, 2002); however, in *C.elegans* PB2 appears to form at the site where PB1 was emitted (Dorn et al., 2010). Furthermore, in *C. elegans* Aurora kinase (AIR-2) localizes to both polar bodies which appear in close proximity (Schumacher et al., 1998). In echinoderms, starfish PB1 is tethered to PB2 and not linked directly to the egg surface (Harada et al., 2003), as is the case in the sea cucumber (Miyazaki et al., 2005). It should be noted that in many invertebrate species including annelids, nemerteans, molluscs and echinoderms that the meiotic spindles are astral and thought to possess centrioles, while the meiotic spindles in *C.elegans* and chordates (including ascidians) are not thought to possess centrioles (Crowder et al., 2015b). There is therefore no correlation between centriole presence and tethered polar bodies. Moreover, even if centrioles were involved in positioning PB2 emission site, the same questions would arise: how does PB1 become tethered to PB2 instead of being directly linked to the egg, and is the midbody remnant found between PB1 and PB2? However, exceptions to PB1 tethering are found, notably in the vertebrates (*Xenopus* and mouse), whereby both PB1 and PB2 are each linked directly to the egg surface (Dehapiot et al., 2013; Li et al., 2016 and Figure 7, scenario 2). We do not know the reason for this difference, however since vertebrate oocytes arrest at Metaphase II (none of the invertebrate species listed in Table 1 arrest at Meta II) for an extended period, it is possible that this long arrest gives the chromosomes on the second meiotic spindle ample time to define the site of PB2 emission using a Ran-GTP gradient only. Furthermore, in Meta II arrested mouse oocytes some evidence indicates that the midbody formed during PB1 emission may be lost, since GFP::Plk1 labels midbody 1 during PB1 emission, but there is no remaining GFP::Plk1 labelling of the midbody remnant in Meta II arrested oocytes (Wianny et al., 1998).

Some aspects of the temporal order of events during PB emission have emerged from elegant studies in *Xenopus* and mouse. Early during meiosis I in *Xenopus*, an actin cap is present before the fall in Cdk1 activity at the end of meiosis I, and it is the fall in Cdk1 that leads to the recruitment of active Cdc42 during anaphase to the actin cap causing outpocketing (Zhang et al., 2008). In *Xenopus* oocytes however, since MPF activity falls to low levels long before Cdc42 cortical recruitment is observed, it has been suggested that anaphase-specific spindle changes (occurring as a consequence of the low MPF activity) act as the precise temporal trigger for Cdc42 cortical recruitment (Zhang et al., 2008). In mouse oocytes, active Cdc42 also forms a cortical cap above one pole of the first meiotic spindle once the spindle has migrated close to the cortex (Wang et al., 2013), and a second cap of active Cdc42 is present during PB2 emission (Dehapiot et al., 2013). Similarly, in the mouse oocyte it had previously been shown that the cortical outpocket of PB1 was abolished by preventing the fall in Cdk1 activity, but not by preventing homologue disjunction (Herbert et al., 2003; Kudo et al., 2006). Thus, in both *Xenopus* and mouse oocytes, the activation of the anaphase promoting complex (APC), which leads to the destruction of cyclin B (inactivating MPF) and securin (activating separase), is permissive for the cortical recruitment of active Cdc42 driving outpocketing during anaphase. Finally, although the fall in MPF activity is permissive for outpocketing, it is not clear how active Cdc42 is recruited to the actin cap during anaphase, although spindle pole proximity to the cortex is thought to be required (Leblanc et al., 2011). Whether chromatin is also involved in driving Cdc42 recruitment to the actin cap during outpocketing (when spindle microtubules are depolymerized) is difficult to assess in mouse oocytes, because microtubule depolymerization activates the spindle assembly checkpoint (SAC) thus preventing the fall in MPF activity (Homer et al., 2005). However, in mouse oocytes DNA beads can induce an actin cap and more importantly a cortical outpocket also forms near the beads (Deng and Li, 2009). It should be noted that although microtubules were not detected around the DNA beads it is possible that some microtubules were not detected in these immunofluorescence images (Deng and Li, 2009). This is similar to our findings using ascidian eggs: we found that following microtubule depolymerization, which disrupts the spindle but does not entirely remove all tubulin staining from around the chromosomes (Figure 6), the female chromosomes can still induce a cortical outpocket that resorbs. It should be noted that ascidian eggs do not possess a SAC, so the MPF activity declines despite a depolymerized spindle (the eggs form a pronucleus at the correct time) thus allowing us to detect the cortical outpocket when MPF activity falls to low levels (Dumollard et al., 2011). Thus, in oocytes it is not entirely known whether chromatin and/or the spindle pole triggers Cdc42 accumulation at the actin cap triggering outpocketing.

Based on the results presented here, we suggest that the midbody directs the precise site where the actin cap forms during meiosis II, and that this cortical site prefigures the precise location of cortical outpocketing during PB2 emission. For example, our evidence indicates that actin is recruited around the midbody early during meiosis II (perhaps in part due to a Ran-GTP gradient coming from the adjacent chromosomes activating Cdc42) at the site where cortical outpocketing will later occur during Anaphase II. We also propose that for efficient PB2 emission one pole of the second meiotic spindle is attracted to the midbody, and that during cortical outpocketing this drives spindle rotation as the free spindle pole is forced to tilt into the egg interior. Interestingly, we noted that prior to tilting, the second meiotic spindle moved laterally along the cortex (see Supp Movie 4). In addition, we propose that the tilting of the spindle moves one spindle pole and its associated chromatids away from the cortex so that only one cortical outpocket is formed rather than two. Indeed, when the spindle fails to rotate two simultaneous cortical outpockets are formed, one on either side of PB1 (Figure 5). However, we still do not know how one pole of the spindle is chosen to be captured by the polarized subdomain of cortex centered on the midbody, nor what triggers the lateral spindle displacement. Interestingly, in *C.elegans* 2-cell stage embryos, the midbody remnant influences astral microtubules during nucleus-centrosomal complex rotation thus biasing the orientation of the mitotic spindle in the P1 cell (Singh and Pohl, 2014). Also, it has recently been demonstrated in preimplantation mouse embryos that the cytokinetic bridge connecting two sister blastomeres acts as a scaffold leading to microtubule stabilization and outgrowth during interphase (Zenker et al., 2017). So microtubules could potentially grow from either the spindle pole or even from the midbody remnant to influence second meiotic spindle position (but we do not discount the involvement of an actin-based mechanism). However, the identity of these microtubules is not known, and indeed in the ascidian egg the meiotic spindle is not thought to possess centrioles or astral microtubules. However, the absence of centrioles does not preclude the presence of short astral microtubules. For example, in *Xenopus* and *C. elegans* oocytes that also lack centrioles, the meiotic spindle poles displays short astral microtubules (Crowder et al., 2015a; Gard, 1992). Unfortunately, due to the density of microtubules in the meiotic spindle it is difficult to detect astral microtubules in the ascidian. It should also be borne in mind that meiotic spindles can rotate in the complete absence of midbodies. For example, the first meiotic spindle rotates during meiosis I independent of the midbody in the ascidian. Nonetheless, since tethered polar bodies are a widespread occurrence throughout the invertebrates, we propose that the site of the previous PB1 cytokinesis directs the precise positioning of PB2 formation (Table 1). Thus, we wonder whether this phenomena of midbody directed PB2 emission is linked to rapid progression though meiosis II in oocytes that do not arrest at Meta II (the vast majority of invertebrates: exceptions are chaetognaths and amphioxus). Meiosis II lasts only 15 min. in the ascidian. In species that do not display tethered polar bodies such as the vertebrates, oocytes can remain arrested at Meta II before fertilization for several hours. Finally, this proposition is somewhat similar to the well-studied case of budding yeast, in which each new daughter cell emerges adjacent to the previous cytokinetic ring (« bud scar ») (Chiou et al., 2017).

Overall, we propose that the midbody remnant that remains within the egg following PB1 emission plays two roles to ensure the spatial precision of meiosis II. First, it likely directs the cortical polarization which defines the location of outpocketing, perhaps via the chromatin-centered Ran-GTP pathway. Second, the midbody remnant and the cortical polarized subdomain appear to attract one spindle pole and cause the second meiotic spindle to tilt. We show here that if the second meiotic spindle does not tilt, then emission of PB2 is perturbed, perhaps because both spindle poles and both sets of chromatids at meiotic Anaphase II come to lie in close proximity to the cortex, causing double polarization of the cortex on either side of PB1. This can lead to failure of efficient PB2 emission as two outpockets can form rather than one outpocket and one PB2. However, it should be noted that in some species of Clam (*Corbicula leana*) failure to rotate the meiotic spindle is a natural event. In *Corbicula leana* two first polar bodies form simultaneously leading to a complete loss of the egg chromosomes thus causing androgenesis of the fertilized egg (Komaru et al., 2000). Finally, we point out the obvious limitation of the current work which is mostly descriptive and correlative due to our inability to either destroy or remove the midbody remnant without also destroying or removing either the actin cap or the egg chromosomes respectively.

## ACKNOWLEDGEMENTS

This work was supported by the Agence National de la Recherche (ANR-12-BSV2-0005-01 to AMcD), an ARC grant (to RD), NSF 12445 (to DB), support from Sorbonne Universities ANR-11-IDEX-0004-02 to the Picard Network (to AMcD) and also an EMBRC-Fr visiting fellowship (to DB). We thank Laurent Giletta for animal collection and Stefania Castagnetti for *Mytilus* gametes. We also thank Sameh Ben Aicha for assistance with MoveInCell (http://movincell.org/).

## AUTHOR CONTRIBUTIONS

A.McD. conceived project and conducted experiments.

A.McD wrote the ms.

J.C, R.D, G.P,V.C, D.B. and C.H. performed experiments.

C.H provided molecular tools.

## Figures and Supplementary Movies (with links)

**Supplementary Movie 1. Bright field movie showing location of protrusion between PB1 and PB2** http://movincell.org/medias/view/id/346

Example of normal PB2 emission in the ascidian *Phallusia mammillata*. Note the small polar corps between PB2 and PB1.

**Supplementary Movie 2. PB2 emission in the bivalve *Mytilus galloprovincialis*** http://movincell.org/medias/view/id/347

Bright field movie showing that PB2 and PB1 are tethered in the bivalve due to the precise emission of PB2 at the previous site of PB1 emission. In *Mytilus* however we do not see a polar corps protruding from the surface of PB2. This is either because there is no protruding polar corps in *Mytilus* as in the ascidian, or alternatively because the protrusion is obscured due to PB1 and PB2 being compressed together in *Mytilus*.

**Supplementary Movie 3. Midbody 1 and midbody 1 location between polar bodies** http://movincell.org/medias/view/id/348

Confocal z stack from a live fertilized egg containing Plk1::Ven. Example of the midbody between PB1 and PB2 (midbody 1: arrow) and between PB2 and the egg (midbody 2: arrow). PB1 and PB2 are indicated by the arrows. The pronucleus is also indicated by an arrow.

**Supplementary Movie 4. Second meiotic spindle tilts during emission of PB2** http://movincell.org/medias/view/id/349

Fluorescence images of Ens::3GFP labelling of microtubules extracted from a confocal 4-D time-lapse sequence showing emission of PB2. Note that the second meiotic spindle tilts between 7 and 11 min: first a rapid translocation moves the second meiotic spindle horizontally along the cortex (from 7 to 8 min.). This translocation ends when one pole of the spindle becomes positioned near the site of PB1 and the spindle simultaneously tilts (starting at 9 min.). PB2 is emitted at 13 min. in this movie (please note that this is actually 25 min. after fertilization). Also note that the microtubules of the large sperm aster are also labelled with Ens::3GFP. Scale bar 40μm. Time in min.

**Supplementary Movie 5. Two second PB2 outpockets** http://movincell.org/medias/view/id/350

Bright field images of a fertilized egg displaying two second cortical outpockets, one on either side of PB1.

**Supplementary Movie 6. Failed rotation of second meiotic spindle displaying two PB2 outpockets** http://movincell.org/medias/view/id/351

Unfertilized eggs were injected with a mixture of Ens::3GFP and Kif2::mCherry mRNA and left overnight to translate protein products then fertilized. Bright field, Ens::3GFP (green), Kif2::mCherry (red) and overlay are shown. The second meiotic spindle fails to rotate and two outpockets form above both sets of chromosomes following Ana II (labelled with Kif2::mCherry: see 24-31 min.). Scale bar = 40μm.

**Supplementary Movie 7**. **DNA can polarize the egg cortex in ascidians as in mouse oocytes** http://movincell.org/medias/view/id/352

Unfertilized eggs were injected with MAP7::GFP to label the microtubules and left overnight to translate protein product. The following day the injected eggs were bathed in Hoechst 33342 (10μg/ml) to label the chromosomes and 2μM nocodazole to depolymerize the microtubules and then fertilized. Upper row of images: bright field images; second row of images: chromosomes blue; third row of images: microtubule green; bottom row of images: overlay of green and blue images. The meiotic spindle microtubules depolymerized and formed a mass around the egg chromosomes (first column of images). An outpocket forms (second column of images). The outpocket resorbs into the egg (last column of images). n=8.

**Supplementary Movie 8. Cortical actin cap forms under PB1** http://movincell.org/medias/view/id/353

Unfertilized eggs were injected with LifeAct protein coupled to mCherry to label actin and fertilized. The movie plays a z-stack of confocal images showing the labelling of actin by LifeAct::mCherry (red). Scale bar = 20μm. n= 8.

**Supplementary Movie 9. Dynamics of cortical actin cap during meiosis II** http://movincell.org/medias/view/id/369

Unfertilized eggs were injected with LifeAct::mCherry protein to label actin and fertilized. Confocal time-lapse experiment during meiosis II showing emission of PB2. Note the accumulation of actin on the cortex of the egg adjacent to PB1. Cortical outpocket of PB2 appears at 6 min. in the movie (but please note that this is equivalent to 25 min. post fertilization). Note that an actin cap is present before the cortical outpocket appears. Note also that the first midbody is visible at (3-6 min.) and also between PB2 and PB1 (10 and 19 min.). Scale bar = 20μm. n=8.

